## Supplementary figures and images for "Activation of JUN in fibroblasts promotes pro-fibrotic programme and modulates protective immunity"

### Supplemental Fig. S1

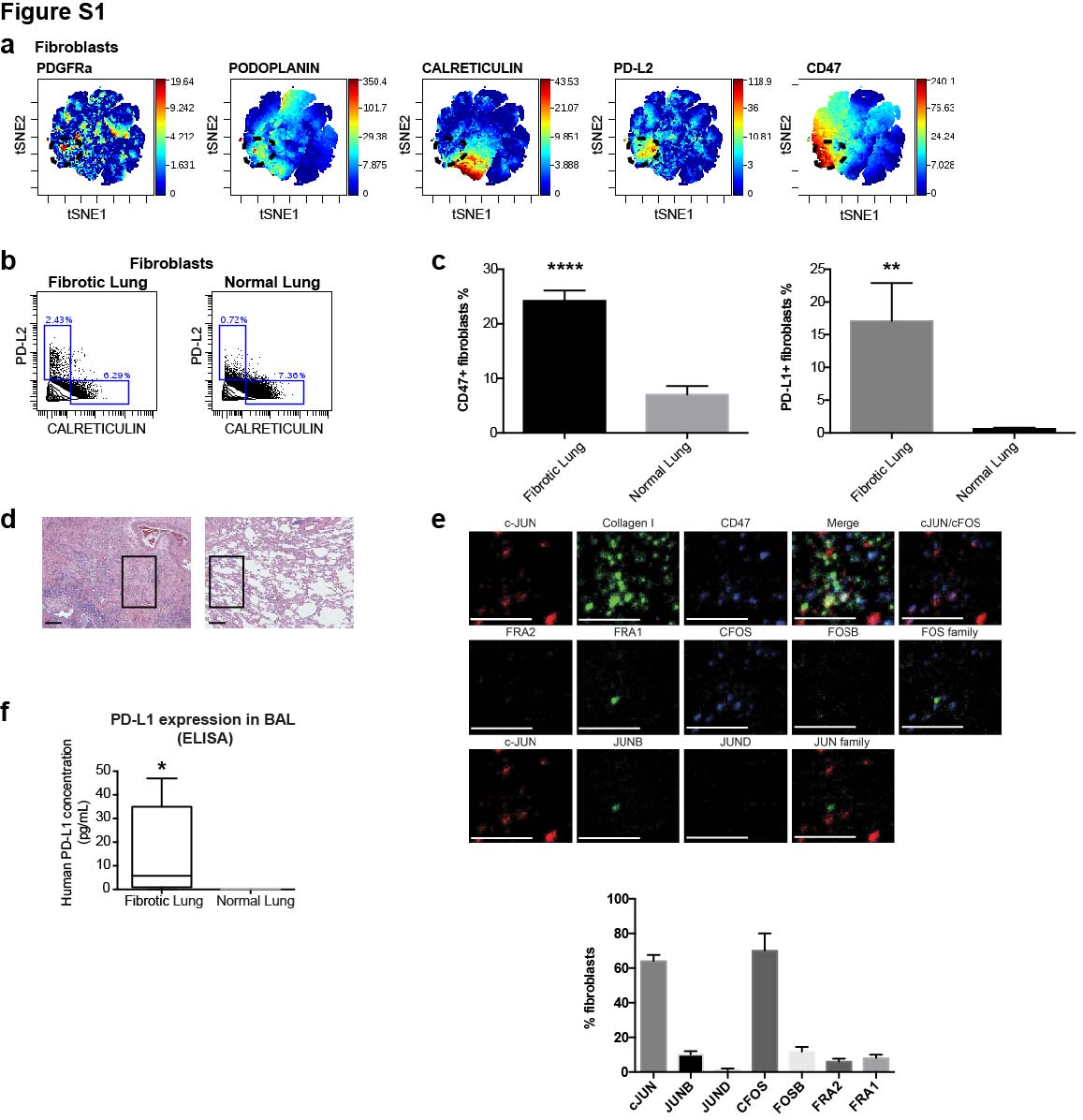

### Supplemental Fig. S2

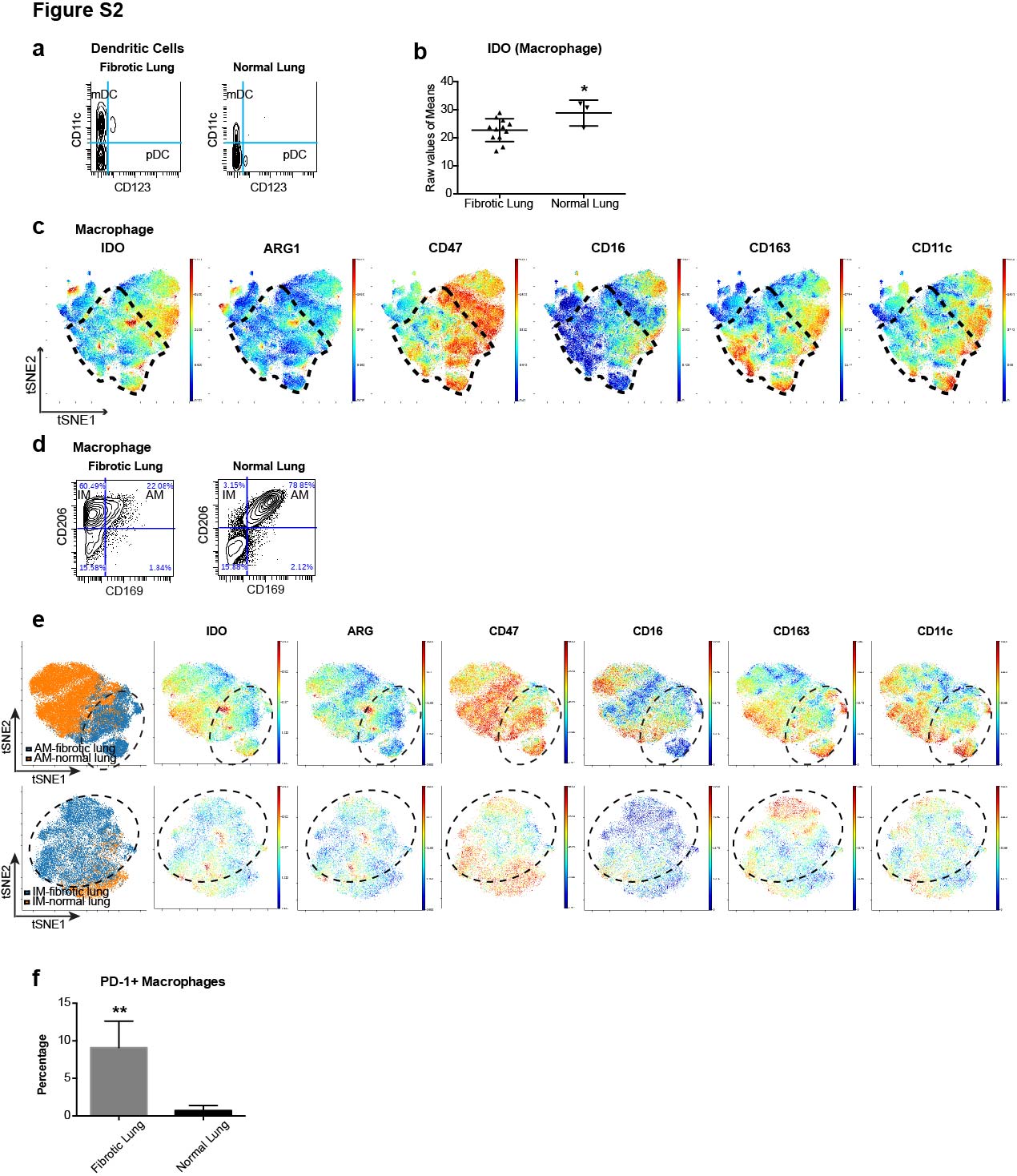

### Supplemental Fig. S3

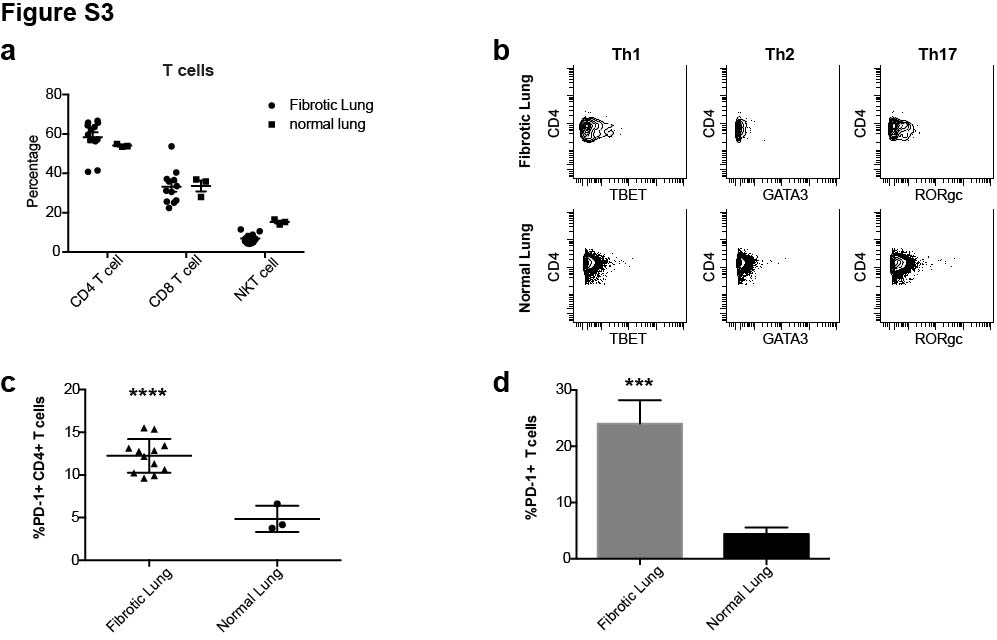

### Supplemental Fig. S4

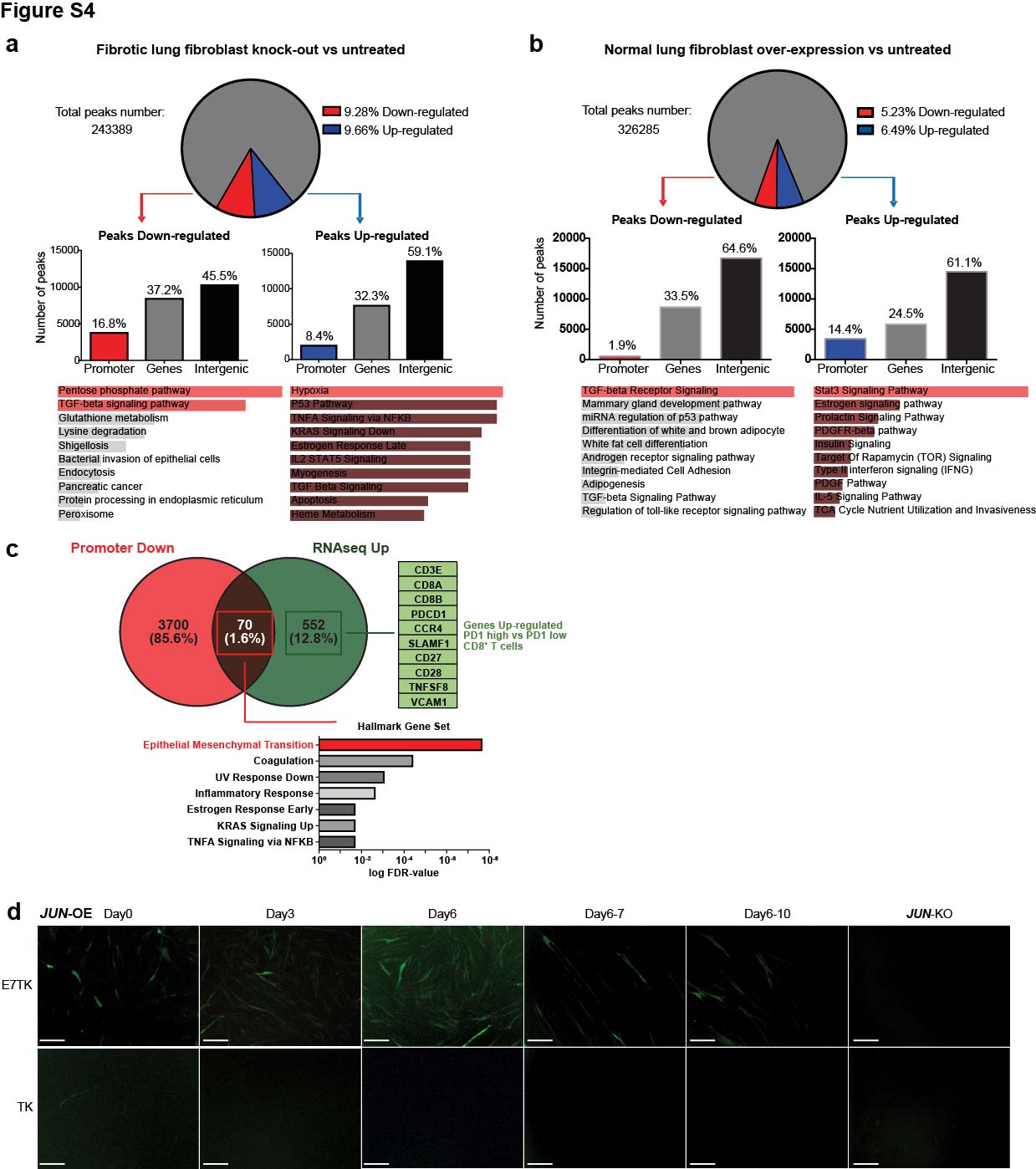

### Supplemental Fig. S5

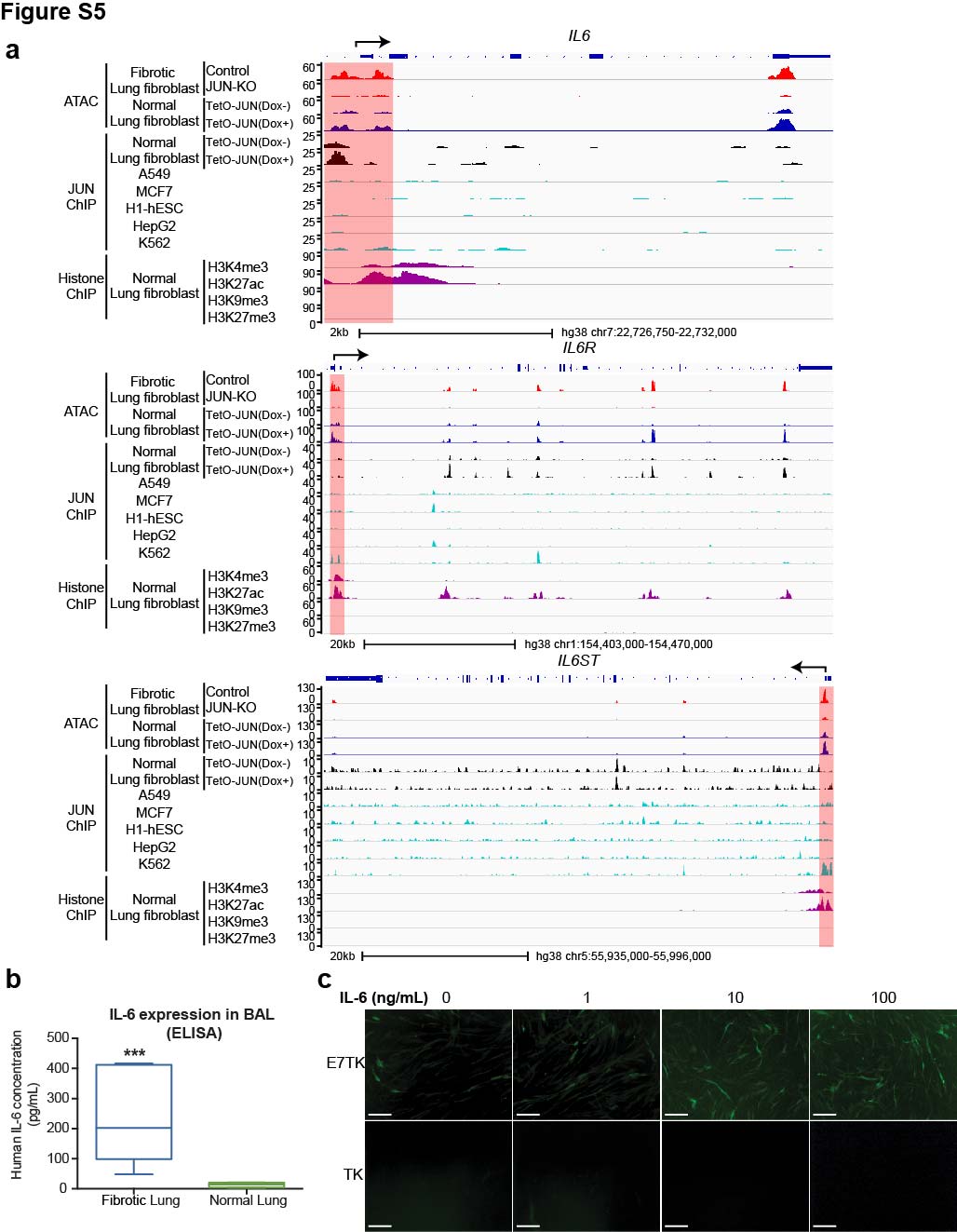

### Supplemental Fig. S6

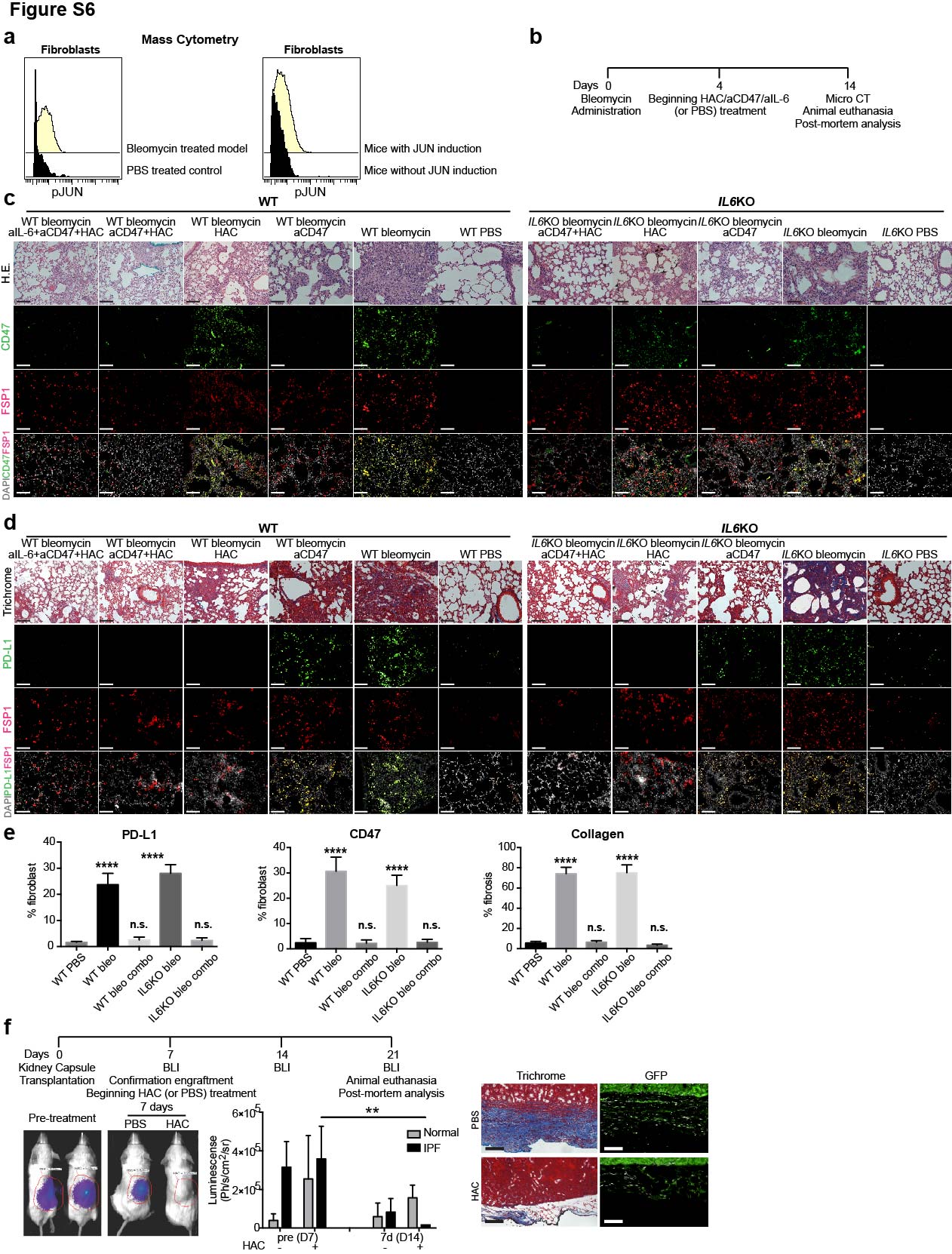

### Supplemental Fig. S7

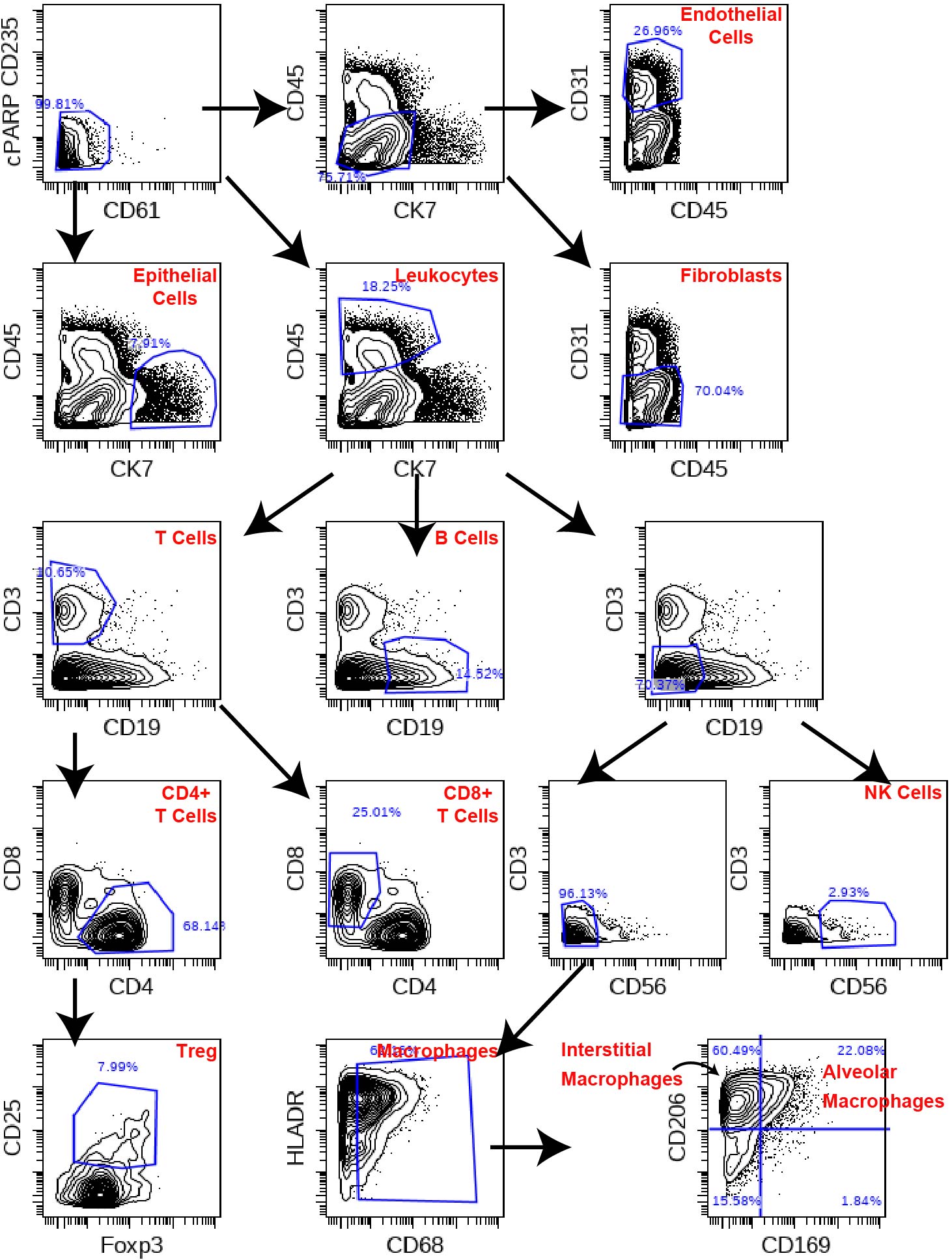
