## Supplemental Table. S1 for "Activation of JUN in fibroblasts promotes pro-fibrotic programme and modulates protective immunity"

**Supplymental Table 1.** Patient infomations.

| Patient # | Age (years) | Gender | Ethnicity | Smoking Status | IPF Stage | Pathology | Oxygen in LPM rest | DLCO % predicted | Comorbidities |
| --- | --- | --- | --- | --- | --- | --- | --- | --- | --- |
| 1 | 61 | Male | Asian | Non-smoker | endstage fibrosing ILD | NSIP, PAH, BAC, honeycomb | 6-8 | 20 | GERD |
| 2 | 69 | Male | Caucasian | Former smoker- | endstage fibrosing ILD | UIP, honeycomb | 6 | 22 | GERD, DM type II |
| 3 | 72 | Male | Caucasian | Former smoker- quit 1976 | endstage fibrosing ILD | UIP, honeycomb | 4-6 | <30 | none |
| 4 | 65 | Female | Caucasian | 2^nd^ had exposure, Non-smoker | endstage fibrosing ILD | UIP, honeycomb, | 4-6 | 25% | Hypercholesterolemia,  DM type II |
| 5 | 69 | Male | Caucasian | Former smoker | endstage fibrosing ILD | UIP, PAH, honeycomb | 8 | <20 | ILD, GERD, depression |
| 6 | 72 | Male | Caucasian | Never smoker | endstage fibrosing ILD | UIP, honeycomb | 8 | 22 | GERD, HTN, Sleep apnea,  ILD, skin cancer, BCC, SCC |
| 7 | 68 | Male | Caucasian | Former smoker- quit 1980 | endstage fibrosing ILD | NSIP | 6 | 20 | SSC, psoriasis, OSA, HTN, obese |
| 8 | 68 | Male | Caucasian | Former smoker- quit 1990, 25 pack years | endstage fibrosing ILD | UIP, honeycomb | 6 | 20 | GERD, HTN, CKD |
| 9 | 63 | Male | Caucasian | Former smoker- quit 1995, 25 pack years | endstage fibrosing ILD | UIP, honeycomb | 10 | 18 | MCI, CAD, HTN, DM type II, hyperlipid, GERD, depression |
| 10 | 54 | Male | Hispanic | Former smoker | endstage fibrosing ILD | chronic hypersens pneumonitis, | 4 | 19 | ILD, GERD, DM type II, TBC |
| 11 | 50 | Female | Asian | Non-smoker | endstage fibrosing ILD | chronic hypersens pneumonitis, PAH | 4 | 23 | GERD, HTN, hyperlipid, DM type II |
| 12 | 59 | Male | Caucasian | Former smoker | normal lung tissue, lobectomy# | normal lung  tumor: lung adenoCA | N/A | N/A | lung adenoCA |
| 13 | 66 | Male | Caucasian | Former smoker | normal lung tissue, lobectomy# | normal lung  tumor: lung adenoCA | N/A | N/A | lung adenoCA |
| 14 | 72 | Male | Caucasian | Former smoker | normal lung tissue, lobectomy# | normal lung  tumor: lung adenoCA | N/A | N/A | lung adenoCA |

*All the ILD patients included in the studies had histologic or radiographic evidence of end-stage fibrosing interstitial lung disease (ILD): UIP (8), fibrotic NSIP (2), fibrotic chronic interstitial pneumonitis (2), DLCO all severe decreased DLCO between <25% of predicted ,FVC <80%, FVC 10% or greater decrement in FVC during 6-month follow-up, 6 minute walk pulse oximetry below 88% or 50m decline in over 6 months, patients **associated pulmonary hypertensive (PAH) features on histopathology. # Our healthy control lung specimens were derived from lung lobectomy specimens for lung cancer, we only received histologic healthy appearing lung distant from the tumor (lung specimen weights 150-200g, tumor diameters ranging 0.8-2 cm, stage pT2pN0).
